# Sticky-flares: real-time tracking of mRNAs… or of endosomes?

**DOI:** 10.1101/029447

**Authors:** David Mason, Raphaël Lévy

## Abstract

The article by Briley *et al* is the latest of several papers based on the use of oligonucleotide-capped gold nanoparticles (or Spherical Nucleic Acids, SNAs) to detect mRNA within cells (1). The technology was developed, published and patented by Prof Chad Mirkin and is commercialised under the trade name SmartFlare. Briefly, the SNAs are hybridised with fluorescent reporter strands (resulting in quenching of the fluorescence by the gold particles) and internalised into cells by endocytosis. The target RNA binds either with the SNA, releasing the reporter strand (SmartFlare) or, in the latest study, with the reporter strand itself, releasing the reporter-strand/target-RNA duplex (Sticky-flares).

Since they enter cells by endocytosis, Sticky-flares (and SmartFlares) need to escape those compartments before they can report on mRNA levels in the cytosol. This essential step is currently not documented in the literature. To the contrary, we (2), as well as the Mirkin group (3), have shown that the overwhelming majority of SNAs remain trapped within endosomes and get degraded leading to an increase of fluorescence. We note that localisation of gold nanoparticles in cells after uptake can be easily determined by electron microscopy (4). In the absence of any mechanism for, or evidence of escape, we suggest that degradation within endosomes is a more likely interpretation for the Briley *et al* results than live imaging of cytosolic mRNAs.

This alternative interpretation is further corroborated by close inspection of the time lapse data which reveals that the moving fluorescent puncta are much larger (~ 0.8 μm) than the diffraction-limited points that would be expected of single fluorophores. Furthermore, the fluorescent spots colocalise with structures which can be clearly seen in the transmitted light images (Figure 1 and movie (5)). Single molecules such as oligonucleotides duplexes cannot be detected in bright field microscopy. Instead the structures and their trafficking appear typical of intracellular vesicles.

**Figure 1:**
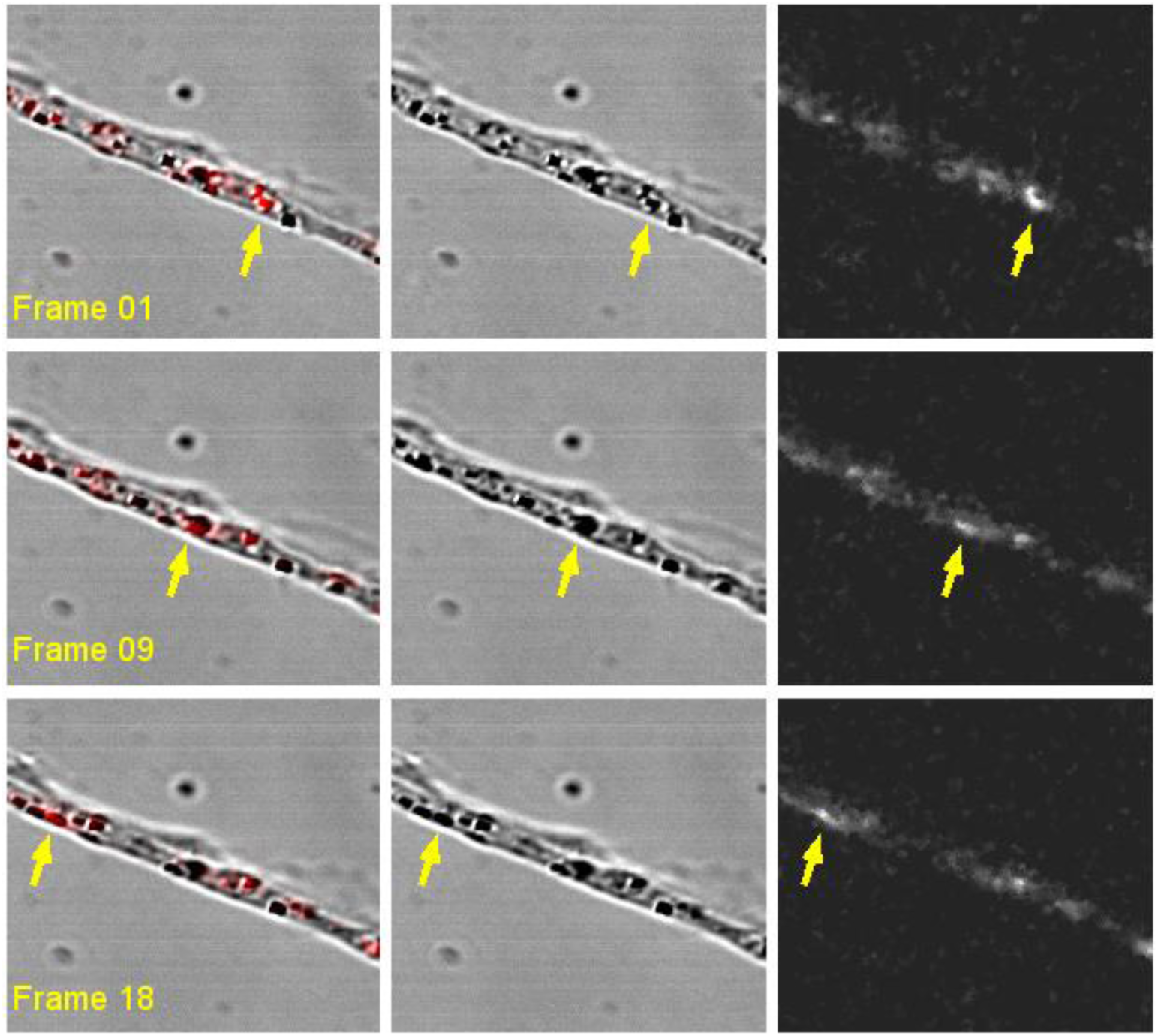
Sticky-flares signal colocalises with intracellular vesicles visible in the bright field channel; we thank Chad Mirkin for providing the raw data of movie S1 of Briley et al that were used to generate this figure. The raw data have been processed in the following ways: a Gaussian band-pass filter (50px) in Fourier space (transmitted channel) to correct uneven illumination; a median filter of 1 pixel in the fluorescent channel to despeckle the image; the image was cropped to just a representative area and enlarged to 150% to aid visualisation. The full movie can be found on FigShare (5).

While it is beyond the scope of this letter to analyse in details each figure in Briley *et al,* we note that care must be taken in the interpretation of selected thin-optical microscopy images and that the lack of an essential control (the effect of the β-actin siRNA on uptake) makes the interpretation of the siRNA data difficult. The development of a new technology requires validation with the current standard. Here we suggest that, in addition to direct evidence of endosomal escape, the authors should perform Fluorescence In Situ Hybridisation (FISH) at the end of a live Sticky-flares experiment and systematically and quantitatively compare the localisation of the fluorescence signal.

